# Computational prediction of protein interactions on single cells by proximity sequencing

**DOI:** 10.1101/2023.07.27.550388

**Authors:** Junjie Xia, Hoang Van Phan, Luke Vistain, Mengjie Chen, Aly A. Khan, Savaş Tay

**Affiliations:** Pritzker School of Molecular Engineering, The University of Chicago, Chicago, IL, 60637, USA; Division of Infectious Disease, University of California, San Francisco, CA, 94143, USA; Lymphocyte Biology Section, Laboratory of Immune Systems Biology, NIAID, NIH, Bethesda, MD, 20892, USA; Section of Genetic Medicine, Department of Medicine, The University of Chicago, Chicago, IL, 60637, USA; Department Human Genetics, The University of Chicago, Chicago, IL, 60637, USA; Department of Pathology, The University of Chicago, Chicago, IL, 60637, USA

## Abstract

Proximity sequencing (Prox-seq) measures gene expression, protein expression, and protein complexes at the single cell level, using information from dual-antibody binding events and a single cell sequencing readout. Prox-seq provides multi-dimensional phenotyping of single cells and was recently used to track the formation of receptor complexes during inflammatory signaling in macrophages and to discover a new interaction between CD9/CD8 proteins on naïve T cells. The distribution of protein abundance affects identification of protein complexes in a complicated manner in dual-binding assays like Prox-seq. These effects are difficult to explore with experiments, yet important for accurate quantification of protein complexes. Here, we introduce a physical model for protein dimer formation on single cells and computationally evaluate several different methods for reducing background noise when quantifying protein complexes. Furthermore, we developed an improved method for analysis of Prox-seq single-cell data, which resulted in more accurate and robust quantification of protein complexes. Finally, our model offers a simple way to investigate the behavior of Prox-seq under various biological conditions and guide users toward selecting the best analysis method for their data.

## Introduction

Advances in single cell sequencing have enabled unprecedented analyses of cellular heterogeneity in complex biological systems^1,2^. Single-cell RNA sequencing^3^ (scRNA-seq) is among the most widely used methods. However, because proteins are the effector molecules for the majority of biological functions, RNA data alone is not sufficient to investigate these protein functions thoroughly. Signaling events, for example, typically begin with receptor clustering, protein phosphorylation, and other protein-protein interactions, all of which occur prior to transcription.

To investigate the roles of protein interactions in greater depth, we recently developed a method called proximity sequencing (Prox-seq) for simultaneous quantification of mRNA, surface proteins and protein complexes at the single-cell level^4^. Prox-seq captures protein complex information in barcoded DNA oligonucleotides (oligos) using a proximity ligation assay^5,6^ (PLA). Each protein in Prox-seq is targeted by two DNA-conjugated antibodies, called Prox-seq probes A and B (Figure 1a). The DNA oligos on probes A and B are ligated only if two protein molecules are sufficiently close to each other. The result of this ligation is referred to as a “PLA product.” The ligation distance is expected to be 50-70nm^7^. In order to generate a PLA product, the oligo belonging to a Prox-seq probe A must ligate to the oligo belonging to a Prox-seq probe B. Importantly, unligated probes do not contribute to the signal because both library preparation and sequence alignment require barcodes from both the A and B probe. Upon sequencing, the number of PLA products can be determined by counting the number of unique molecular identifiers (UMIs). Because of this design, the number of PLA products measured for a protein is a reflection of both the abundance of that protein and the availability of nearby Prox-seq probes. By combining Prox-seq with scRNA-seq, these PLA products can be sequenced alongside complementary DNA (cDNA) libraries, providing information on gene expression, protein abundances, and protein complex formation from single cells^4^.

**Figure 1.**
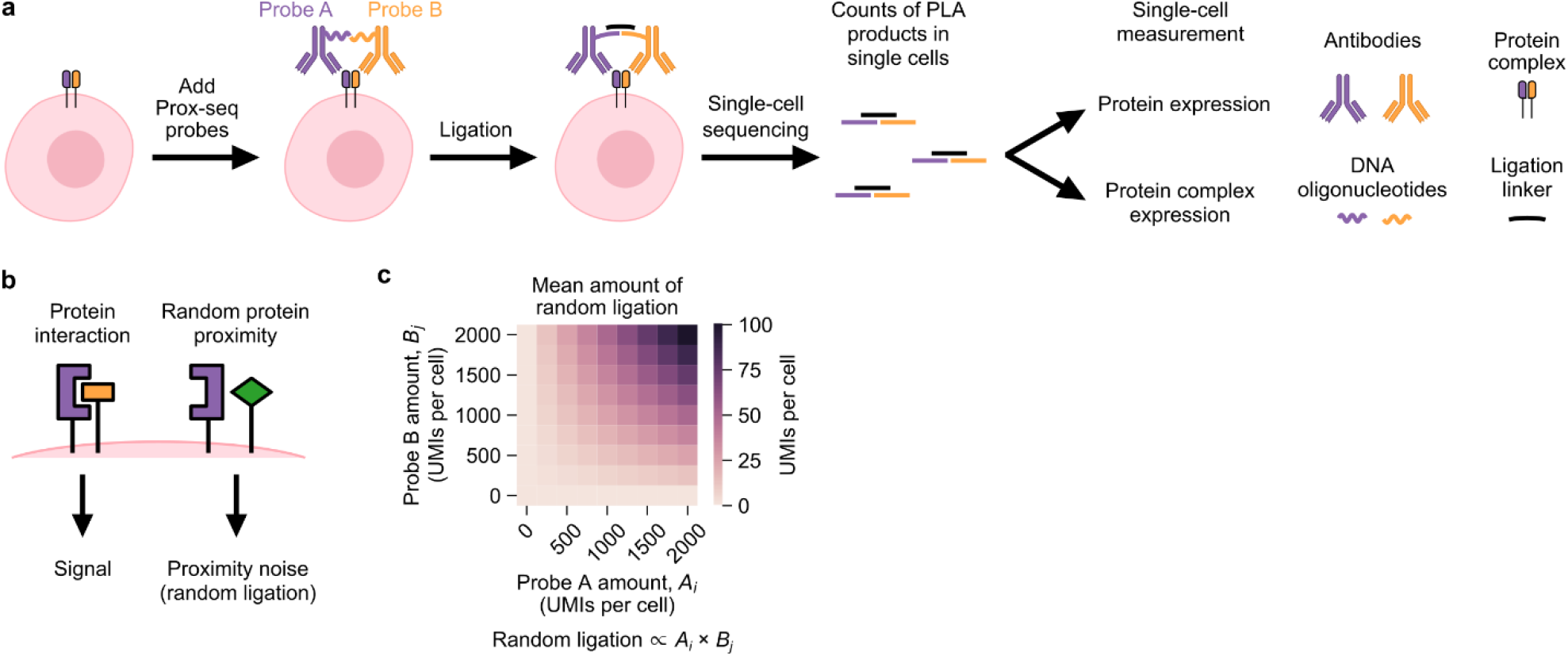
Working principle of Prox-seq and identification of proximity noise. (a) Schematic showing the main steps of Prox-seq. (b) Schematic showing the background in Prox-seq that is caused by proximity noise (random ligation of non-interacting protein molecules). (c) Heatmap showing the expected amount of proximity noise created from simulations of two protein molecules at varying expression levels. By modeling the mean amount of proximity noise with a binomial distribution (see Methods), we found that it was proportional to the product of the abundances of the two protein molecules.

Prox-seq protein data contains a unique source of background noise, namely the ligation of two protein molecules that do not functionally interact but are nevertheless sufficiently close to each other by random chance. We call this effect “proximity noise” (Figure 1b). Proximity noise exists because the average distance between probes on the cell surface decreases with increasing protein abundance (see Methods). A previous study showed that proximity noise led to false positive detection of protein interactions for in situ PLA^8^. A theoretical model showed that the mean amount of proximity noise is proportional to the product of the expression levels of the two proteins that made up the PLA product (Figure 1c). In short, the presence of PLA products for a specific pair of proteins does not guarantee that the two proteins functionally interact and form stable complexes.

To account proximity noise, we previously proposed and used a statistical method, termed the iterative method, to differentiate protein complexes from random ligation in PLA product counts^4^. Initially, this method establishes an “expected value” for each PLA product, representing the number of PLA products that would exist if Prox-seq probes were randomly distributed across the cell surface. Subsequently, the method subtracts the expected background from the PLA product counts. If a PLA product’s count exceeds its expected value, the difference between observed and expected PLA products is attributed to non-random protein complexes. This procedure is iteratively executed for every type of PLA product in each individual cell. Although this method successfully recovered positive controls of known protein complexes, assessing its performance on experimental data is challenging, as the generation of new Prox-seq datasets often lacks comprehensive knowledge of the entire set of protein complexes and their expression levels.

In this study, we present a simulation model for single-cell proteomic data in proximity sequencing experiments and use it to computationally benchmark the performance of several new and existing protein complex prediction methods. After calibrating the model with experimental data, the simulation model allowed us to quantitatively analyze proximity noise and its effects on the measured PLA product counts. We compare the performance of three methods: the iterative method, a new linear regression-based method, and a new ensemble method that combines the two. We find that, while the iterative and linear regression-based methods perform well in several different scenarios, combining them into a single method yielded the most accurate and robust quantification of protein complexes. These results shed insight onto how the spatial organization of surface proteins translate into Prox-seq data and provides guidelines for use of Prox-seq and related dual-binding technologies for multi-omic analysis of single cells.

## Results

### Overview of the simulation model

Based on a physical model of how PLA products are formed in each single cell, we created a simulation model of PLA product count data. We reasoned that proximity alone would determine if a Prox-seq probe A and a Prox-seq probe B ligate and produce a PLA product. We constructed the simulation model in a way that allowed us to simulate probes that bind to non-interacting protein molecules (proteins that are not part of a complex) separately from probes that bind to interacting molecules (proteins that are part of a complex). This procedure enabled us to independently tune the abundance of proteins and protein complexes in the simulation, and to observe how these properties affected Prox-seq data.

First, we generated the non-interacting Prox-seq probes A as random points on a sphere (Figure 2a). These points indicate that the protein molecules exist as monomers; thus, any PLA products they form would be caused by proximity noise and as a result of being in a protein complex. Further, we assumed such protein monomers were distributed randomly on the cell surface. Then, we repeated the process to generate the non-interacting Prox-seq probe B signal. Second, we generated the interacting Prox-seq probes A and B by generating a sphere of random points. These points corresponded to detectable protein complexes. Because these two probes A and B both bound to the same protein complexes, the Prox-seq probe A points would necessarily be in proximity with their corresponding Prox-seq probe B points. Finally, any pairs of probe A and B with Euclidean distances less than the ligation distance were considered ligated and produced PLA products (see Methods). If a probe A was within the ligation distance with more than one probes B, one such probe B was chosen at random to ligate with said probe A.

**Figure 2.**
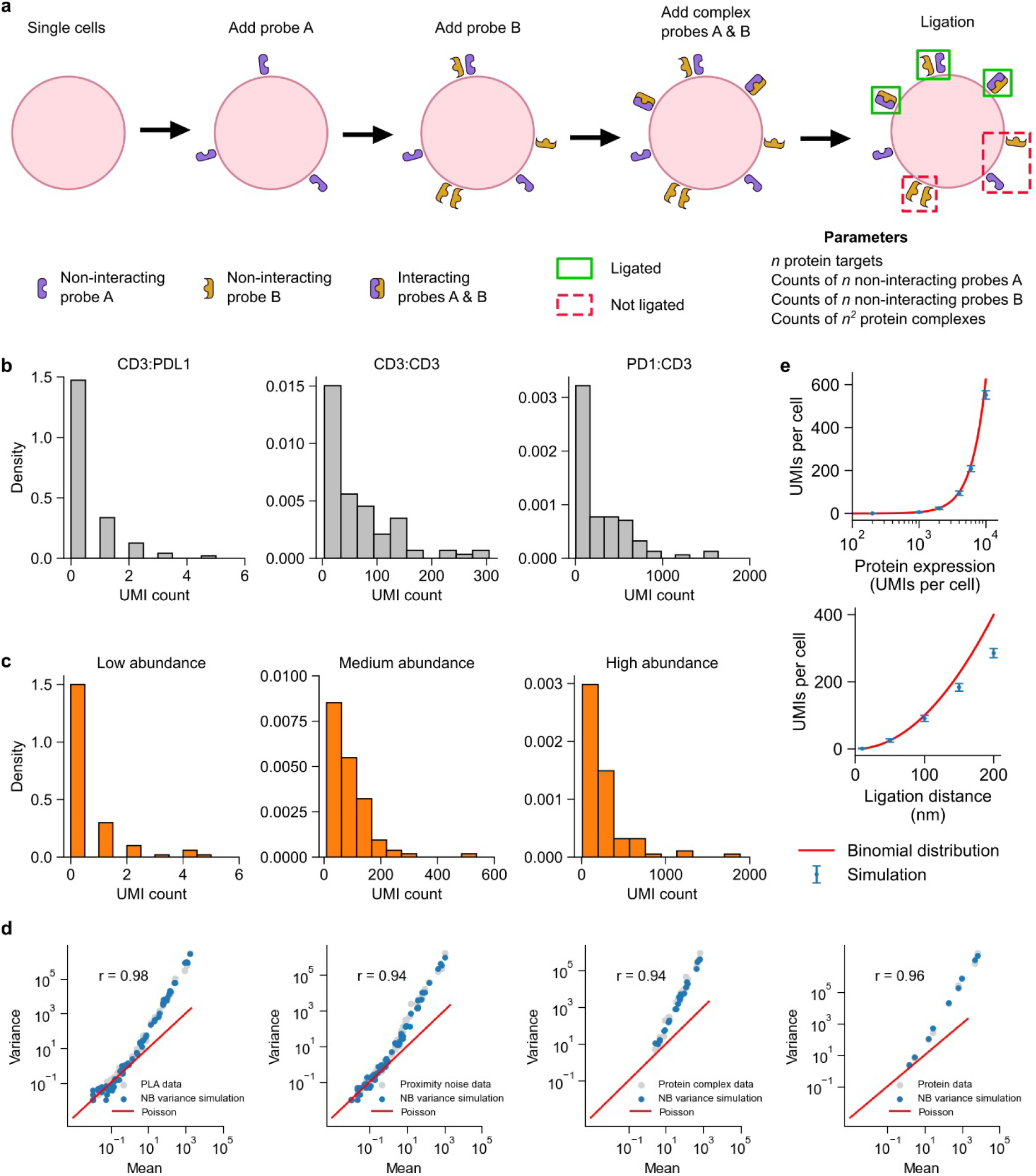
Overview and calibration of simulated Prox-seq data. (a) Schematic for the simulation model of PLA products. The simulation was separately performed on a cell-by-cell basis. First, a number of non-interacting probes A and non-interacting probes B were added as random points on a sphere. Next, a number of protein complexes were added as random points on a sphere. These points corresponded to probes A and B that bound to interacting protein molecules. Finally, probes A and B that had a Euclidean distance lower than the ligation distance were ligated, thus creating PLA products. (b) Histograms showing the UMI counts of three example PLA products in single Jurkat cells. (c) Histograms showing the UMI counts of three example simulated PLA products with NB variance. (d) Scatter plots of mean-variance relationship show how negative binomial variance captures overdispersion in PLA data, proximity noise data, protein complex data, and protein data. (e) The relationship between proximity noise (measured as UMI’s) and protein abundance (top) or ligation distance (bottom).

We next compared the simulated Prox-seq data to experimental data. We analyzed T cells (Jurkat cell line) and B cells (Raji cell line) with a panel of Prox-seq probes that targeted both T cell and B cell markers from a previously reported study^4^. Simulated counts of PLA product and protein expression followed the Poisson distribution, whereas the experimental data exhibited overdispersion (Figure S1a, S2a). We found that adding variance in the form of a negative binomial distribution (NB) for non-interacting probes and protein complexes was sufficient to capture the overdispersion of the real data (NB variance, see Methods). With the added NB variance, the simulated data, like the experimental data, had a right-skewed distribution across different PLA product abundances (Figure 2b, c). Notably, the simulation model with added variance captured the positive correlation between observed PLA product count and non-proximal probe count in real data (Figure S1b-g). The simulation model with no variance, however, showed a negative correlation between PLA product count and non-proximal probe count (Figure S1d and S1e). The NB variance model also produced non-proximal probe counts with similar distributions to those observed in experimental data (Figure S2).

We generated replicated datasets by sampling from the fitted model for posterior predictive checks (PPCs)^9^. We then assessed how well these data samplings maintained the properties of the observed data with two metrics. First, we measured the similarity between the coefficient of variation per PLA product, proximity noise, protein complex and protein. This comparison enables evaluation of how well the mean-variance relationship of real data is preserved (Figure 2d & Figure S3a). Second, we perform Mann-Whitney U-test statistic to measure the extent to which the replicated data and raw data come from the same distribution (Figure S3b). Finally, we characterized the amount of proximity noise in the most basic scenario when there were no protein complexes detectable by the Prox-seq probe panel. The simulation demonstrated that the amount of PLA product produced by random ligation scales quadratically with both protein abundance and ligation distance (Figure 2e). These results show that our model and simulations faithfully capture key aspects of real Prox-seq data in single cells and reiterates the importance of identifying and removing proximity noise, which can especially be large for highly expressed proteins.

### Iterative prediction of protein complex abundance

An iterative method was used to previously identify the existence of stable protein complexes in Prox-seq measurements. This method proposed that when there were no protein complexes, the observed count of a PLA product i:j could be calculated from the abundance of the probe A targeting protein i, and the probe B targeting protein j (see Methods). This calculation resulted in an expected random count for PLA products that represents the PLA count caused by proximity noise. We reasoned that if the observed count of PLA product i:j was higher than the calculated expected random count, then i:j indicated a non-random protein interaction. To quantify the protein complexes on each single cell, we calculated the difference between the observed and expected random PLA product count (Figure 3a). This method was called the iterative method, because it involved solving a system of quadratic equations (describing all possible protein dimers) iteratively (see Methods)^4^. This method relied on the fact that Prox-seq can measure protein abundance, similar to flow cytometry and CITE-seq^10^. The abundance of a protein was the amount of protein molecules that were present on the cell surface, and therefore included both molecules in monomeric and complex forms. In our previous study^4^, we proposed that the protein abundance could be estimated from Prox-seq data by summing the appearances of each protein across its associated PLA products (see Methods). Here, we find by using our simulated data that such an estimate is a good approximation of the true protein abundance, as they are strongly correlated (Figure S4).

**Figure 3.**
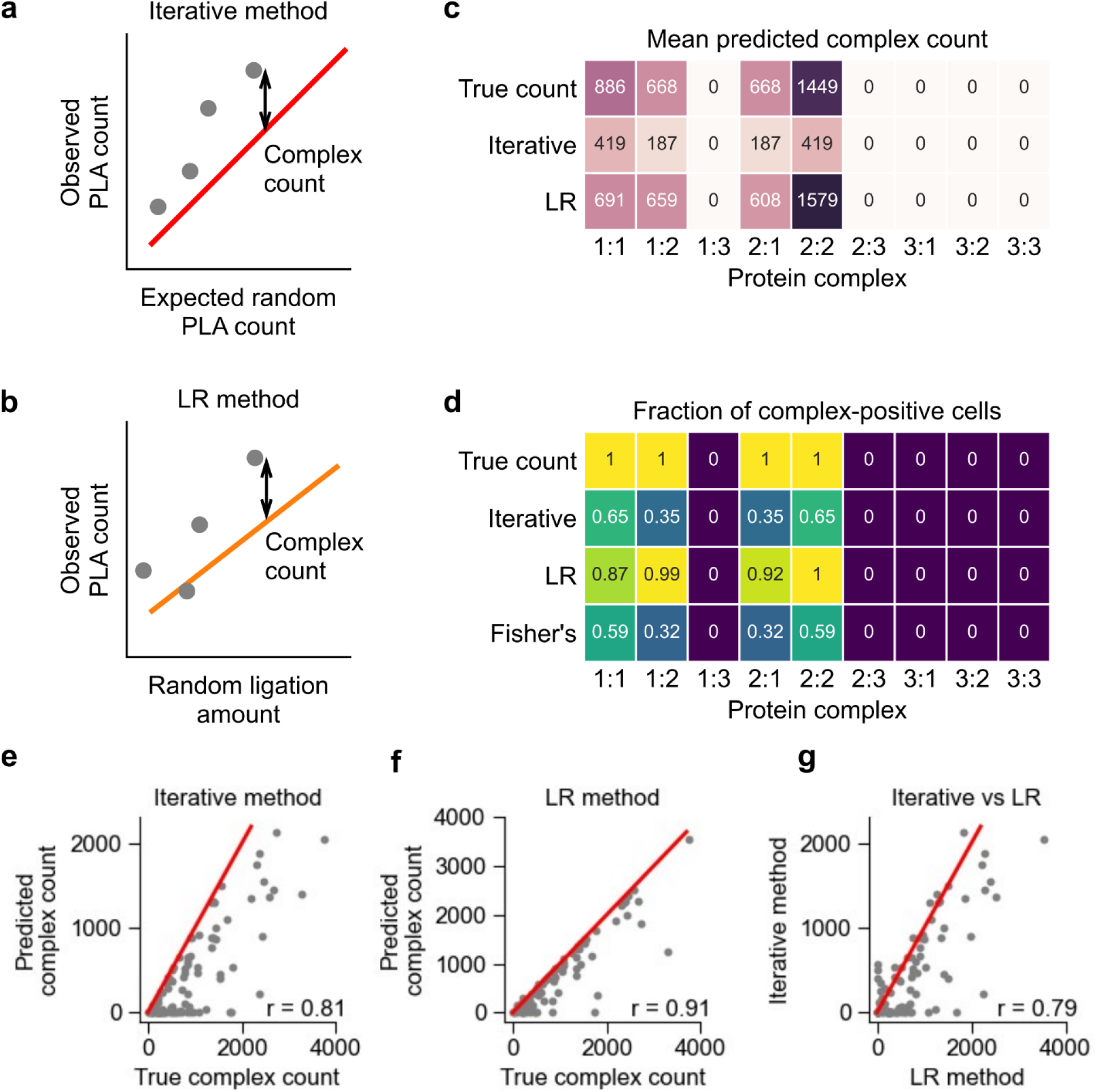
Comparison between the iterative and linear regression (LR) methods for protein complex prediction in simulated data. (a, b) Schematics showing the working principle of (a) the iterative method and (b) the LR method. In the iterative method, the protein complex count is the difference between the observed and expected PLA product count. In the LR method, the protein complex count is the difference between the observed PLA product count and its expected amount of random ligation, which is calculated from the non-proximal probe count. In (a), the red line indicates y = x. In (b), the orange line indicates the linear regression fit. (c) Heatmap showing the mean complex count of simulated data, and of the iterative and LR methods’ prediction results. (d) Heatmap showing the fraction of cells expressing a protein complex, as predicted by the iterative method, the LR method, and Fisher’s exact test. In (c, d), the true count represents the ground truth of protein complex count in the simulation. (e, f) Scatter plots showing the simulated and predicted count of protein complex 1:1 using € the iterative and (f) the LR method. (g) Scatter plot comparing the predicted count of protein complex 1:1 from the iterative and the LR methods. In (e-g), the red lines indicate y = x, and each dot represents a single cell.

To further examine the assumptions underlying the iterative method, we now constructed the following simulation scenario: The simulation had three protein targets, called protein 1, protein 2 and protein 3. These proteins did not interact with themselves, nor with any other proteins. Furthermore, protein 3 had a lower non-interacting probe count (mean of 100 UMIs/cell compared to 1000 UMIs/cell for proteins 1 and 2, Table S1). Simulated data showed that our assumptions behind the iterative method were correct. When there were no interactions between the proteins, the observed PLA product counts were similar to the expected random count (Figure S5a). When we introduced the protein complex 1:1 to the simulation while keeping the other parameters the same, the observed counts of the PLA product 1:1 was higher than its expected random count (Figure S5b).

One weakness of the iterative method is complexity of hyper-parameter tuning, which can result in sub-optimal convergence. They key parameter is the initialization setting, which are the initial estimates of protein complex abundances. By default, the algorithm assigns and initial value of 0 to all protein complexes. However, different initialization settings will influence iterative behaviors to convergence, as well as tolerance (Figure S6a). Unsensible initialization tends to generate nonsensical predictive outputs. (Figure S6b). This led us to consider more robust methods for protein complex quantification.

### Prediction of protein complex abundance using linear regression

To address the weakness of the iterative method we developed a new approach (the linear regression - LR method). This method uses an experimentally modified Prox-seq procedure that enables direct measurement of Prox-seq probes that were not ligated because they were not proximal to another Prox-seq probe (we refer to these as non-proximal probes)^4^. The proximity noise for a PLA product i:j should be proportional to the product of the non-proximal probe A targeting protein i, and the non-proximal probe B targeting protein j. We reasoned that if linear regression is used to model the observed PLA product count onto the estimated random ligation amount, true protein complexes would have positive intercepts (see Methods). The slope was then used to estimate the amount of random ligation, and the count of a protein complex was calculated by subtracting the estimated random ligation from the observed PLA product count (Figure 3b). Experimentally, we observed strong heteroscedasticity in the PLA product count when regressed on to the random ligation amount (Figure S7). Therefore, we performed linear regression using weighted least squares instead of ordinary least squares (see Methods).

We created a new simulation to directly compare the iterative and LR methods. The simulation’s parameters were set to approximate the experimental data. More specifically, the simulation had three protein targets: protein 1, protein 2 and protein 3. Proteins 1 and 2 interacted both with themselves and each other (Figure 3c, Table S1). Protein 3 did not interact with itself, nor with protein 1 or protein 2. Furthermore, protein 3 had very low non-interacting protein count (mean of 2 UMIs/cell compared to 20 and 15 for proteins 1 and 2, respectively). We found that the iterative method correctly identified protein complexes 1:1, 1:2, 2:1 and 2:2 (Figure 3c).

To determine if we can statistically infer the enrichment of PLA products, we performed a one-sided Fisher’s exact test on the counts of PLA products (Figure 3d, see Methods). This analysis correctly identified the four protein complexes present in the sample, independently confirming that the generated protein complexes occur at a higher frequency than random and can be statistically inferred (Figure 3d, see Methods). With regards to quantification of protein complexes on single cells, we observed that the iterative method consistently underestimated the true protein complex count (Figure 3c, e). Conversely, the LR method not only correctly identified the four true protein complexes (complexes 1:1, 1:2, 2:1 and 2:2), but also produced much more accurate counts for them (Figure 3c, d, f). Overall, the results of the two methods were correlated on the single-cell level (Figure 3g).

### Ensemble method that combines both the LR and iterative methods for analysis of Prox-seq data

We next chose to explore a method that had the potential to outperform both the LR and iterative methods. As shown previously, the major weakness of the iterative method is its sensitivity to initialization conditions. We reasoned that the output from the LR method could be used as a sensible initialization for the iterative method (Figure 4a). Starting the iteration close to the correct result would make it less likely that the method would fall into a spurious local optimization. The performance of all three methods was compared in two simulations: one in which a high percentage of proteins were in complex with other proteins (high signal) and one in which a low percentage of proteins were in complex (low signal) (Table S1).

**Figure 4.**
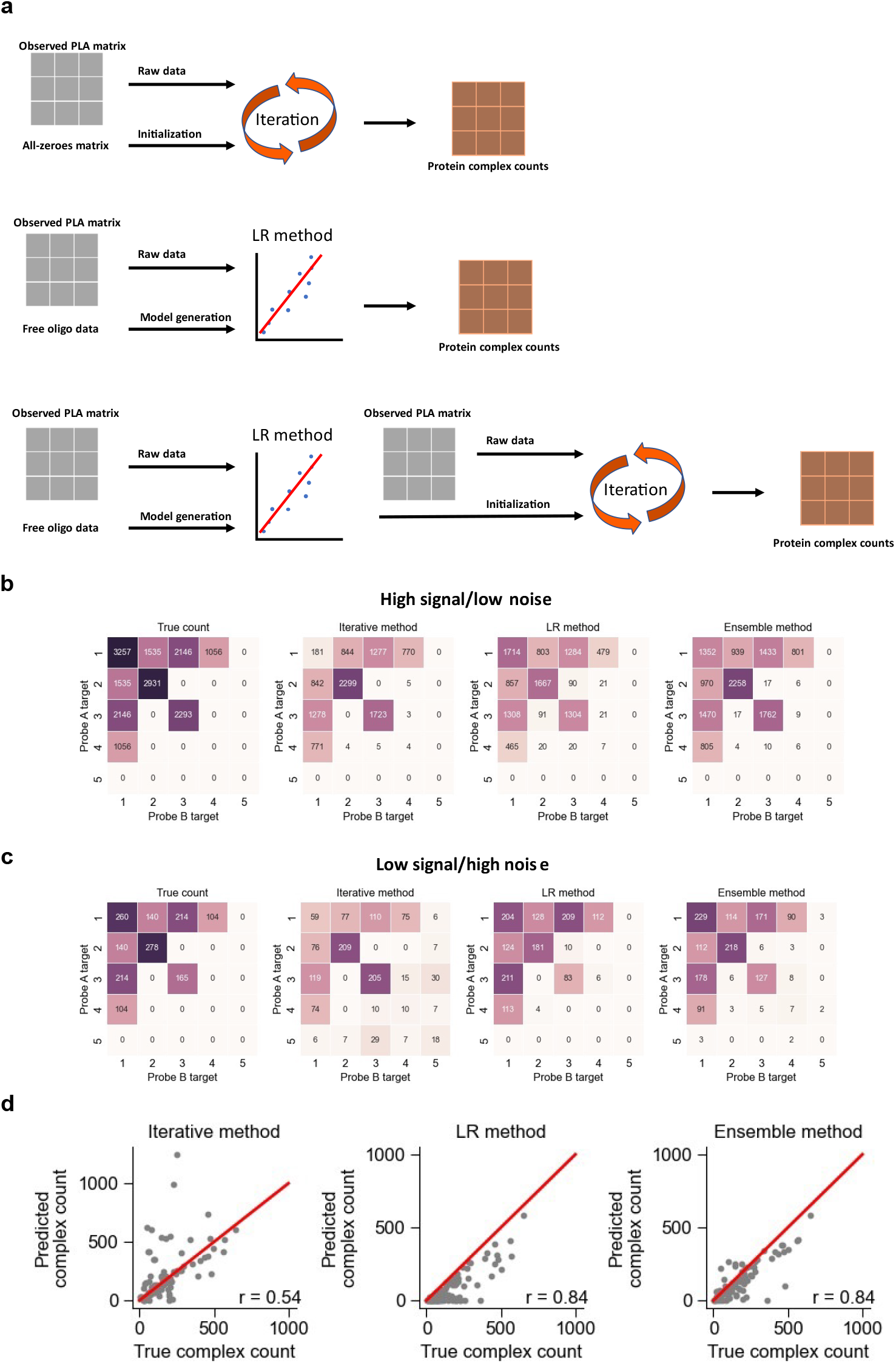
The ensemble method for improved analysis of Prox-seq data. We combine the iterative and LR methods for better prediction of protein complexes. (a) Schematic showing how all three methods arrive at protein complex estimation. The iterative method combines raw data and an initialization with an all-zeroes matrix to quantify protein complexes. The LR method uses raw data and free-oligo data to construct a linear regression model that quantifies protein complexes. The ensemble method begins with applying the LR workflow and uses the output of it to initialize the iterative method. (b) Comparison of all three methods in a regime of high signal and low noise, compared to the true counts. (c) Comparison of all three methods in a regime of low signal and high noise, compared to the true counts. (d) The Pearson’s correlation between true counts and the outputs for each method across single cells. Each example shows complex 3:3 from the low signal/high noise regime.

The iterative method performed well when signal was high, but generated false positives when signal was low (Figure 4b-c). The LR method performed better in the high noise simulation but suffered from false positives when noise was low (Figure 4c). This is not surprising because LR method depends on performing regression with product of non-proximal probes as the explanatory variable and if there are few non-proximal probes across all single cells (or low noise in our simulation), LR method will become unstable. Both methods consistently underestimated the abundance of protein complexes. For the iterative method, this is partly because expected PLA count we assumed is the maximal proximity noise it might have. The slope we use to quantify protein complexes from LR method tends to be larger than the true value because counts of non-proximal probes we can measure are inevitably lower than real counts both in experiment and simulation, which would give us a smaller positive intercept and protein complex count. In contrast, the ensemble method was able to maintain strong performance in both scenarios. It was less likely to produce a false positive, assigned fewer reads to false positives than other methods, and was closer to the true count for most of the protein complexes (Figure 4b-c). Finally, for a given PLA product, the ensemble method was more accurate in quantifying the abundance of true-positive complexes in single cells (Figure 4d).

### A quantitative scoring strategy to comprehensively evaluate prediction methods

To evaluate the predictive performance of these methods more comprehensively, we further propose a quantitative scoring strategy to assign a prediction score for every prediction (Figure S8a). We simulate different biological scenarios with our model and score the overall prediction performance of each method by considering sum of absolute deviation between mean true counts and predicted counts (∑ *Mean*_*deviation*_), sum of Pearson correlation coefficient (∑ *Pearson*) across singles cells (Figure S8b), and sum of ratios of false positive prediction (∑ *FPrate*) across single cells (Figure S8c) (see Methods). Comparing the methods across all scenarios showed that the ensemble method had the highest average prediction score and the lowest variance (Figure S8d & Table S2). The ensemble approach effectively improves iterative method and LR method’s generalization to different biological scenarios.

### Comparison of all three analytical methods to real data and performance evaluations

Next, we evaluated the concordance between all three methods on experimental data from single Jurkat and Raji cells. Overall, we found that each method largely agreed on which PLA products were predicted to be protein complexes (Figures 5a-f). While the bulk measurements of protein complexes showed good agreement between methods, the three methods had varying levels of correlation for single cells (Figure 5g, h). In addition, we observed all three method, along with the Fisher’s Exact test, largely identified the same protein complexes (Figure S9).

**Figure 5.**
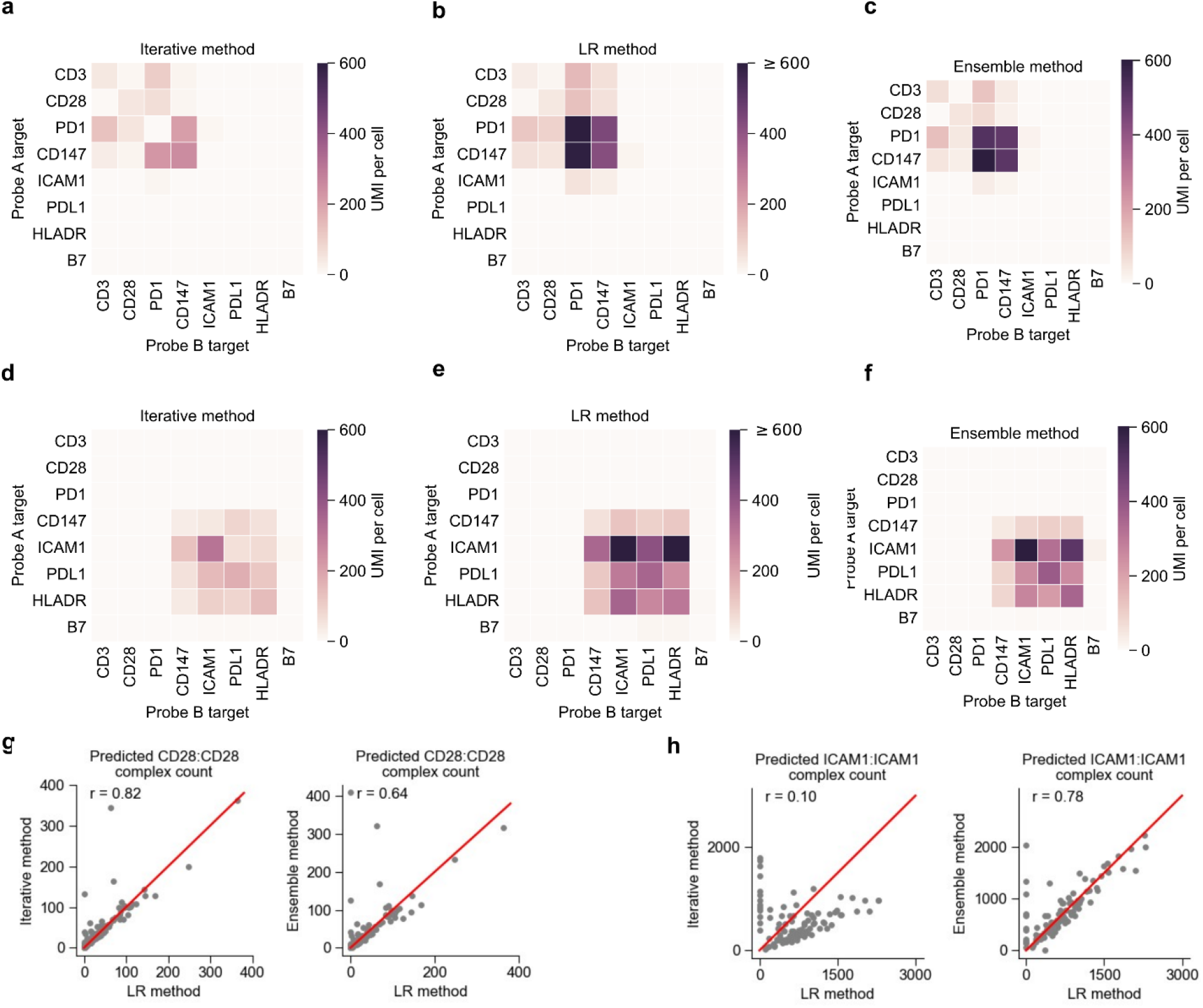
Comparison between the iterative and LR methods on experimental data. (a-c) Heatmaps showing the average of protein complex count, predicted by (a) the iterative method, (b) the LR method, and (c) the ensemble method in Jurkat cells. (d-f) Heatmaps showing the average of protein complex count, predicted by (a) the iterative method, (b) the LR method, and (c) the ensemble method in Raji cells. (g) Comparison of methods for predicting counts of protein complexes of CD28:CD28 and in Jurkat cells. (h) Comparison of methods for predicting counts of protein complexes of ICAM1:ICAM1 and in Raji cells. In (g, h), the red lines indicate y = x, and r indicates the Pearson’s correlation coefficient.

All methods predicted protein complexes CD3:CD3 and CD28:CD28 in Jurkat cells, both of which are known protein complexes^11,12^. All three methods also predicted protein complex ICAM1:ICAM1 in Raji cells, which was shown to dimerize on the cell surface^13^. We also evaluated our methods against a simulation designed to more closely represent the experimental data. Protein expression levels were estimated from the experimental data and used to create simulation models for Jurkat and Raji cells (Table S1). Then, protein complexes corresponding to CD3:CD3, CD28:CD28, and CD3:CD28 were added to Jurkats, whereas HLADA:HLADR and PDL1:PDL1 were added to Rajis. Once again, we observe largely similar performance for all methods (Figure S10).

## Discussion

Here, we presented a comprehensive computational framework for simulating Prox-seq data, and for predicting protein complex count from Prox-seq data. We studied how the quantification of protein complexes was affected by proximity noise, which is caused by proteins that are not functionally interacting but are sufficiently close to each other by random chance to produce valid ligation products. Our simulation model showed that the amount of proximity noise is strongly depended on the protein abundance. Similar results have been observed in commercial *in situ* PLA^8^.

We showed that with respect to protein complex prediction, the iterative method, LR method, and ensemble method largely agree on real experimental data. Therefore, we propose that each of these methods could be used for protein complex detection and quantification, and any protein complexes that were predicted by these methods were highly likely to be true protein complexes. However, in head-to-head comparisons using simulated data, the ensemble method performed well over a larger range of data types than the other methods.

Our simulation model had some limitations. First, it did not consider interactions higher than dimers, diffusion of the protein molecules, their physical sizes, and the technical variability of the Prox-seq assay. Second, the simulation model requires the user to independently select the abundance of a protein complex and its constituents’ non-interacting counterpart. In real cells, these abundances are likely highly correlated. Finally, it assumed that the protein complexes and the non-interacting proteins were uniformly distributed on the cell surface. Despite these limitations, we showed that the overall structure of simulated Prox-seq data is very similar to real Prox-seq data.

Currently, application of each method requires a relatively homogeneous population of single cells. In practice, this requires that simultaneously acquired mRNA data is first used to cluster cell types, and then either method can be applied to individual clusters. This requirement is because each method relied on a statistic of the whole population (the difference between observed and expected random PLA product count for the iterative method, and the linear regression’s intercept and slope coefficient for the LR method) and having different complex expression levels would lower the power the methods. Further study is required to extend these methods to a population of heterogeneous cell types without the use of mRNA data.

We envision that the Ensemble method will be particularly useful when Prox-seq is extended to intracellular proteins. Indeed, since non-specific antibody binding is much more severe in intracellular staining than extracellular staining, random ligation is an even more important source of noise given common macromolecular crowding effect within cells. The simulation model can also be further extended to model Prox-seq data of intracellular proteins. In short, we have validated the protein complex prediction algorithm that was proposed previously^4^, proposed two additional independent methods for protein complex prediction, and introduced a model for simulating Prox-seq data.

## Methods

### Theoretical calculation of proximity noise

Suppose there are A_i_ probes A and B_j_ probes B on the cell surface. Assume that the probes are random points on a spherical surface, and proteins i and j do not interact. Because the ligation distance is significantly shorter than the cell’s radius, we assume that a probe A and a probe B can be ligated if and only if the Euclidean distance between them, *L*, is less than or equal to the ligation distance, *d*_*ligation*_. The Euclidean distance *L* between any pair of random points has the following probability distribution^14^:

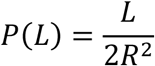

where *R* is the cell radius.

Then, the probability of ligation between two random points on the cell surface is:

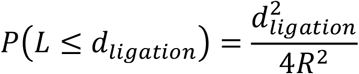

Assume that each probe could be ligated as many times as possible, the mean counts of ligated PLA product i:j, X_i,j_, follow a binomial distribution:

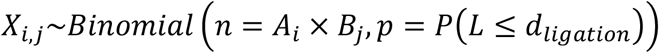

The expected count of PLA product that is created from random ligation of non-interacting probes is:

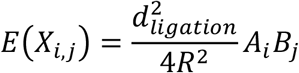

Note that this approximation assumes that each probe can be ligated many times, while the simulation model assumes that each probe can only be ligated at most once.

### Simulation model

Assume that each protein molecule and the Prox-seq probe that binds to it are point particles. Let there be n protein targets. Let A_1_, A_2_,…, A_n_ be the simulation parameters that represent the count of probe A that targets proteins 1, 2,…, n. Let B_1_, B_2_,…, B_n_ be the simulation parameters that represent the count of probe B that targets proteins 1, 2,…, n. Let c_1,1_, c_1,2_,…, c_1,n_, c_2,1_, c_2,2_,…, c_n,n_ be the simulation parameters that represent the counts of protein complexes 1:1, 1:2,…, 1:n, 2:1, 2:2,…, n:n.

The simulation is performed separately on each single cell. For the single cell t, we first generate 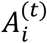 number of random points on a sphere surface, which correspond to the number of detected probe A that targets protein i on cell t. The coordinates of each point are^15^:

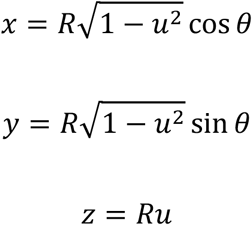

where *R* is the radius of the sphere (taken to be 5 µm, or 5000 units, in our study), *u* is uniformly distributed over [-1,1), and *θ* is uniformly distributed over [0,2π).

Without added variance, 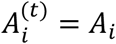. With added negative binomial variance:

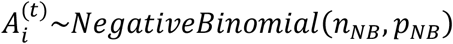

where n_NB_ =1.5 in our study, and 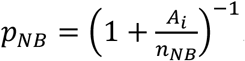. The negative binomial distribution formulated this way provides the probability of getting 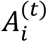 failures, given n_NB_ successes and p_NB_ is the probability of success. n_NB_ is used to control the variance of the probe count, and pNB is calculated such that the mean of 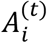 is equal to A_i_.

Second, we randomly generate 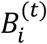 number of points on a surface of a sphere, which correspond to the number of detected probe B that targets protein i on cell t. The coordinates of each point are generated identically to above.

Without added variance, 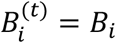. With added variance:

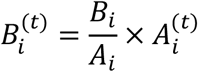

This is to ensure that the counts of detected probe A and probe B that target the same protein are proportional to each other.

Third, we randomly generate 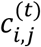 number of points on a surface of a sphere, which correspond to the count of protein complex i:j on cell t. Then, these 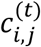 points are added to the previously generated probe A points targeting protein i 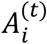, and also to the previously generated probe B targeting protein j 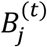.

Without added variance, 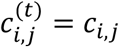. With added variance:

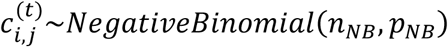

where n_NB_ =1.5 in our study, and 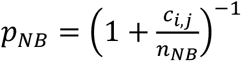.

Fourth, we calculated the pairwise Euclidean distances between all generated probe A points and all generated probe B points. Finally, we randomly go through the pairs of points that are within a ligation distance threshold (chosen to be 50 nm, or 50 units, in our study), and add the corresponding PLA product to the simulated count matrix. Any probe A and probe B points that are chosen are excluded from future PLA products. In other words, each probe A and each probe B can only be ligated at most once.

The number of probe A and probe B points that are not ligated are returned as the simulated non-proximal probe count that is measured by the free oligo modification.

The simulation is repeated 100 times to simulate PLA product counts of 100 single cells. The parameters for all simulations are listed in Table S1. All simulations include negative binomial variance, unless stated otherwise.

### Calculation of protein count and expected PLA product count

The count of a protein i in a single cell is equal to the total number of detected PLA products that are related to the protein i:

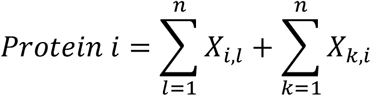

where X_i,l_ and X_k,i_ indicate the observed (i.e., measured) counts of PLA products i:l and k:i, respectively. The PLA product i:i is counted twice towards the protein count to account for the fact that two molecules are present in a homodimer.

The expected count of a PLA product i:j, E_i,j_, is:

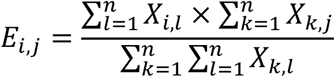

### Protein complex prediction: iterative method

The count of protein complex i:j is calculated iteratively using the following equation:

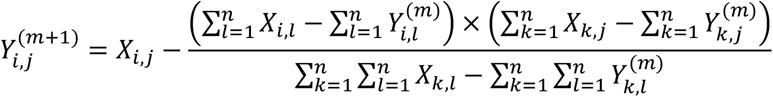

where 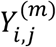 is the predicted count of protein complex i:j at the m^th^ iteration. The initial values for all protein complexes are 0.

The second term of the right hand side represents the count of PLA product i:j that is caused by random ligation.

After each iteration, a one-sided t-test is performed on the values of 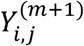 across all single cells. The alternative hypothesis is that the mean of 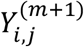 is greater than 1. Next, any 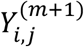 with Benjamini-Hochberg-corrected P-values above 0.05 are set to 0. In other words, any such PLA products were determined to not represent true protein interactions.

There is also a symmetry condition, such that if i:j is a protein complex, then j:i should also be a protein complex, even if 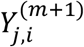 fails the t-test. This is done by setting 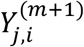 as a fraction of 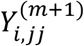:

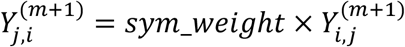

where sym_weight is arbitrarily chosen to be 1 in our study.

### Protein complex prediction: linear regression (LR) method

For each PLA product i:j, its observed count is regressed onto the product of its corresponding non-proximal probe A count and non-proximal probe B count, using weighted least squares:

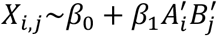

where 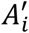 and 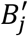 are the count of non-proximal probe A targeting protein i, and non-proximal probe B targeting protein j, respectively. The weight for a sample (ie, a single cell) *p* is:

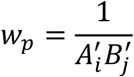

For simulated data, we also scale the interaction term by 10^6^ whenever necessary, such that it is close to the orders of magnitude of X_i,j_. 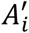 and 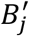 are obtained from PLA products that contain the added free oligos. For example, the count of non-proximal CD3 probe A is equal to the count of PLA product CD3:free_oligo_B, and the count of non-proximal CD28 probe B is equal to the count of PLA product free_oligo_A:CD28.

Next, we performed a one-sided t-test on the intercept coefficient, and the alternative hypothesis is that *β*_0_ > *β*_*cutoff*_. For simulated data, *β*_*cutoff*_ = 1. For experimental data *β*_*cutoff*_ = 10. All PLA products with Benjamini-Hochberg-corrected P-values below 0.05 are considered to be true protein complexes. The protein complex count, Y_i,j_, is calculated as the difference between the observed PLA product count and the interaction term:

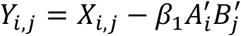

The LR method is related to the binomial approximation of the random ligation signal above. If the counts of non-proximal probes are perfect proxies for the count of non-interacting probes, then we have the following relationship:

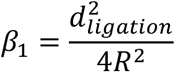

### Protein complex prediction: Ensemble method

The ensemble method relies on solving same quadratic equations as iterative method to approximate counts of protein complex. The only difference is that it takes protein complex matrix calculated from LR method as initial values. There is an argument called df_guess embedded in predictive function which is set to be all zeros by default. Note that LR method should be applied in advance in order to perform ensemble method.

### Protein complex prediction: Fisher’s exact test

For each PLA product i:j, we construct a 2×2 table:

| | Probe B = j | Probe B ≠ j |
| --- | --- | --- |
| Probe A = i | $X_{i,j}$ | $\sum_{\substack{l=1 \\ l \neq j}}^n X_{i,l}$ |
| Probe A ≠ i | $\sum_{\substack{k=1 \\ k \neq i}}^n X_{k,j}$ | $\sum_{\substack{k=1 \\ k \neq i}}^n \sum_{\substack{l=1 \\ l \neq j}}^n X_{k,l}$ |

We perform a one-sided Fisher’s exact test using the table. The alternative hypothesis is that X_i,j_ is higher than the expected value. We perform a Benjamini-Hochberg correction for each cell on the P-values of all PLA products calculated for that cell. The cell is considered to be displaying the protein complex if the corrected P-value is below 0.05.

### Prediction score mechanism

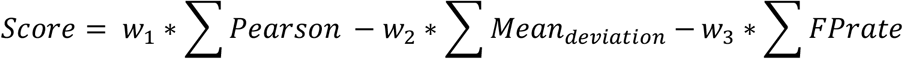

where *w*_1_, *w*_2_, *w*_3_ are chosen to be 0.5, 0.4, 0.1 in our study. ∑ *Pearson* equals to the sum of Pearson correlation coefficients for every real protein complex between true complex counts and predicted complex counts across all singles cells. ∑ *Mean*_*deviation*_ equals to the sum of absolute difference between mean true counts and predicted counts for every PLA product:

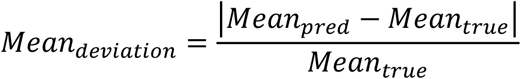

∑ *FPrate* equals to the sum of ratios of false positive prediction across single cells for every non-existing PLA product.

For quantification accuracy evaluation where there are true protein complex counts, we consider parameters ∑ *Pearson* and ∑ *Mean*_*deviation*_. Pearson correlation coefficient takes single cells into consideration while means counts can give us information about bulk abundance of different PLA products. We found that poor prediction of PLA counts in single cells might still contribute to seemingly good mean counts estimation, which shed lights on us that Pearson correlation should be a more important and robust parameter than mean counts. For ∑ *FPrate* evaluation where there is no true complex, we use fraction of complex-positive cells to represent how many ratios of single cells are wrongly assigned at least a complex count. According to our multiple tests, each method tends to assign only few false positive reads, mostly only one in some single cells to PLA products. So that we assume false positive rate a minor metric to be considered in our scoring strategy. In conclusion, we arbitrarily choose effector weight for each parameter given relative importance discussed above.

### Software implementation

All code is implemented in Python3/Anaconda3 (v4.10.3). The code is deposited at https://github.com/tay-lab/Prox-seq_computation.

## Data availability

The raw sequencing data and processed PLA product count data are deposited in NCBI’s Gene Expression Omnibus (accession number GSE196130).

## Acknowledgements

S.T. was awarded an NIH R01 grant GM127527, NIH MIRA/R35 grant R35GM148231, and a Paul G. Allen Distinguished Investigator Award, which supported this work. M.C. was awarded NIH R01 grants GM126553 and HG011883, and an NSF grant 2016307 which supported this work. This work was supported in part by the Intramural Research program of NIAID, NIH.

## Author Contributions Statement

L.V., H.V.P., J. X. and S.T. conceived of and designed the project. H.V.P., J. X., M.C., and A. K. performed statistical and computational analysis. L.V., H.V.P., J. X. and S.T. wrote the manuscript. S.T. supervised the project. All authors reviewed the manuscript.

