## Supplementary File for "Computational prediction of protein interactions on single cells by proximity sequencing"

### Supplementary figures

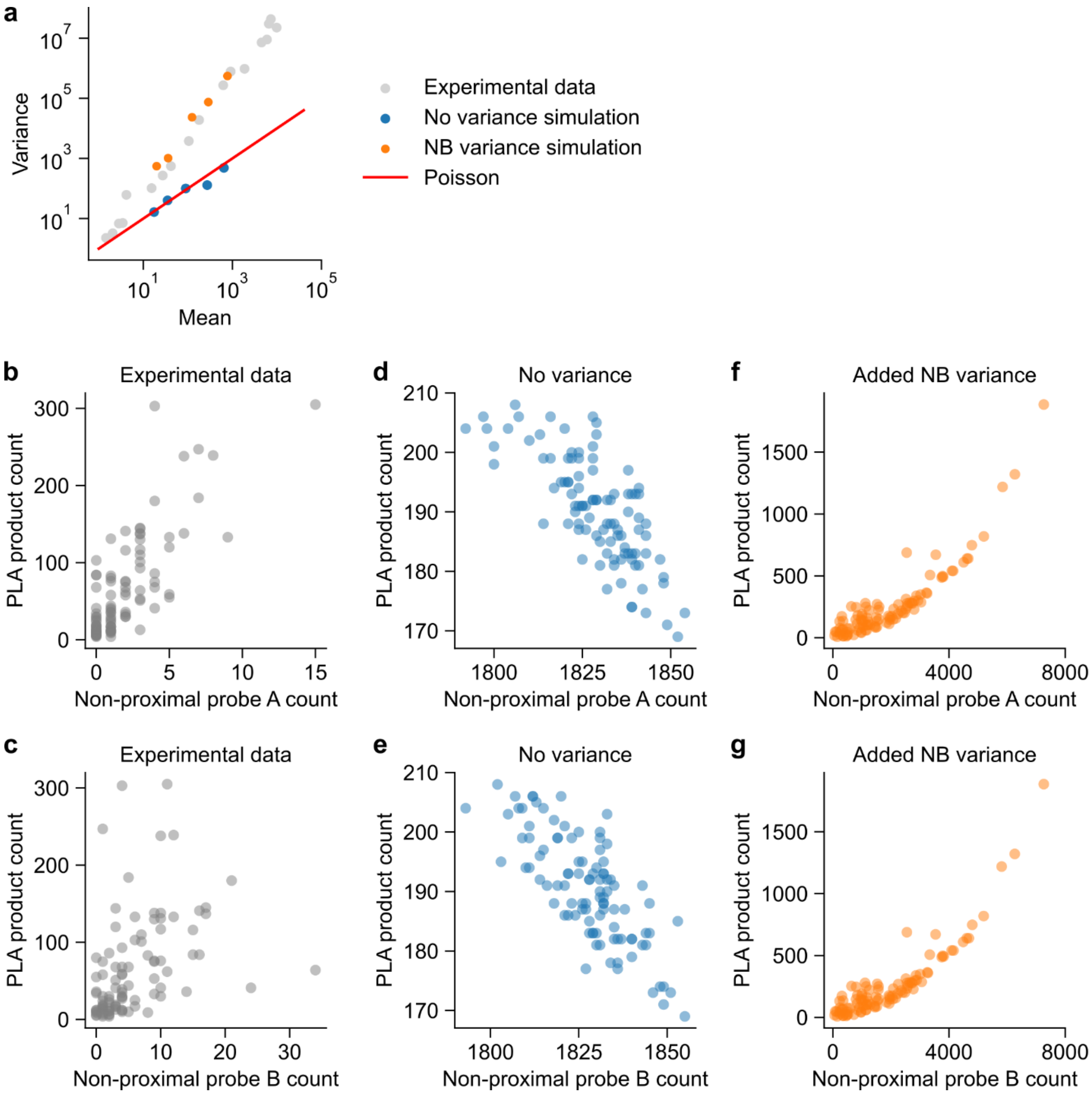


**Figure S1. Comparison of real and simulated data for protein count and non-proximal probe count.** (a) Scatter plot showing the mean-variance relationship in real and simulated protein count. (b, c) Scatter plots showing the relationship between observed CD3:CD3 PLA product and (b) non-proximal CD3 probe A or (c) non-proximal CD3 probe B in Jurkat cells. (d, e) Scatter plots showing the relationship between observed 1:1 PLA product and (d) non-proximal protein 1 probe A or (e) non-proximal protein 1 probe B in simulated data without variance. (f, g) Scatter plots showing the relationship between observed 1:1 PLA product and (f) non-proximal protein 1 probe A or (g) non-proximal protein 1 probe B in simulated data with negative binomial variance.

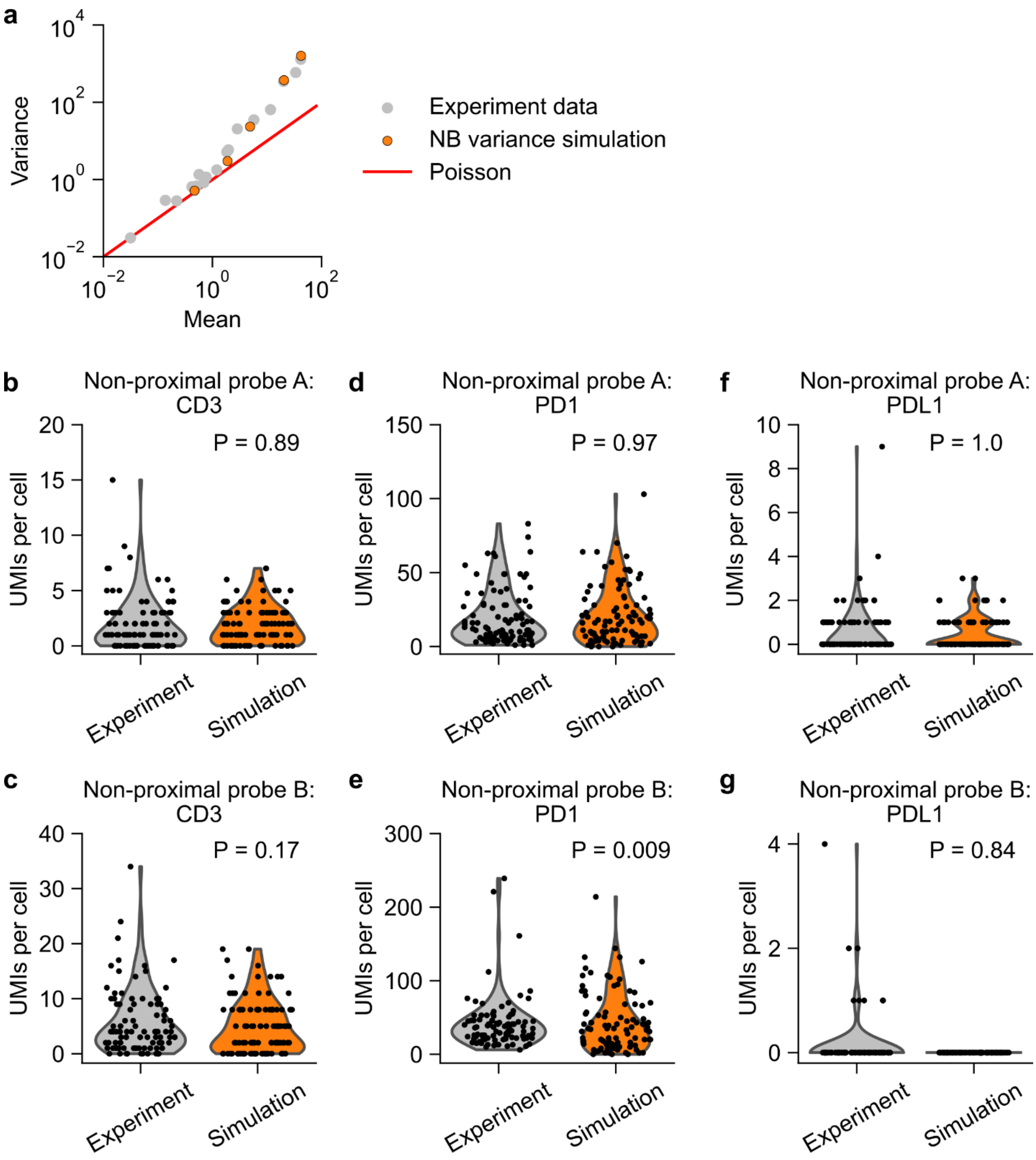


**Figure S2. Distribution of non-proximal probe counts in Jurkat cells and simulated data.** (a) Scatter plot showing the mean-variance relationship in experimental (Jurkat cells) and simulated non-proximal probe count. (b, c) Violin plots showing the experimental and simulated count of (b) non-proximal probe A and (c) non-proximal probe B for CD3 protein. (d, e) Violin plots showing the experimental and simulated count of (d) non-proximal probe A and (e) non-proximal probe B for PD1 protein. (f, g) Violin plots showing the experimental and simulated count of (f) non-proximal probe A and (g) non-proximal probe B for PDL1 protein. non-proximal probe counts for CD3, PD1 and PDL1 proteins were simulated by using the mean non-proximal probe counts in experimental data. Note that Jurkat cells expressed CD3 and PD1 proteins, but not PDL1 protein. P-values are calculated using KS test.

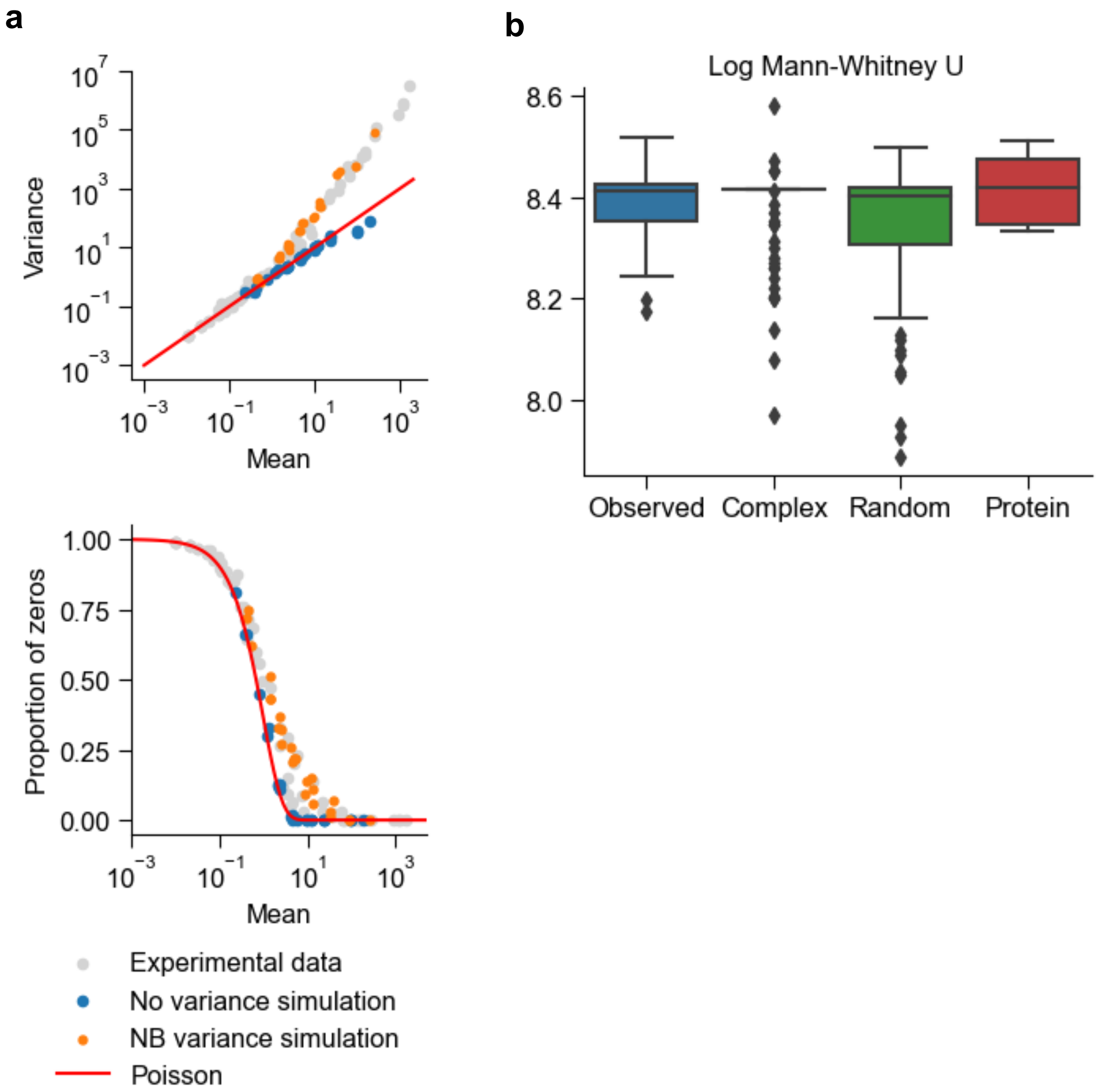


**Figure S3. Negative binomial variance causes the simulation to produce distributions that closely match real data** (a) The simulation with NB variance outperforms Poisson variance for both the mean-variance relationship (top) and proportion of zeroes (bottom). (b) For each observed PLA, protein complex, proximity noise and protein, the Mann-Whitney U statistic between posterior predictive samples and observed data averaged over samples. Box plots indicate the median (center line), interquartile range (hinges), and whiskers at 1.5x interquartile range. Higher is better.

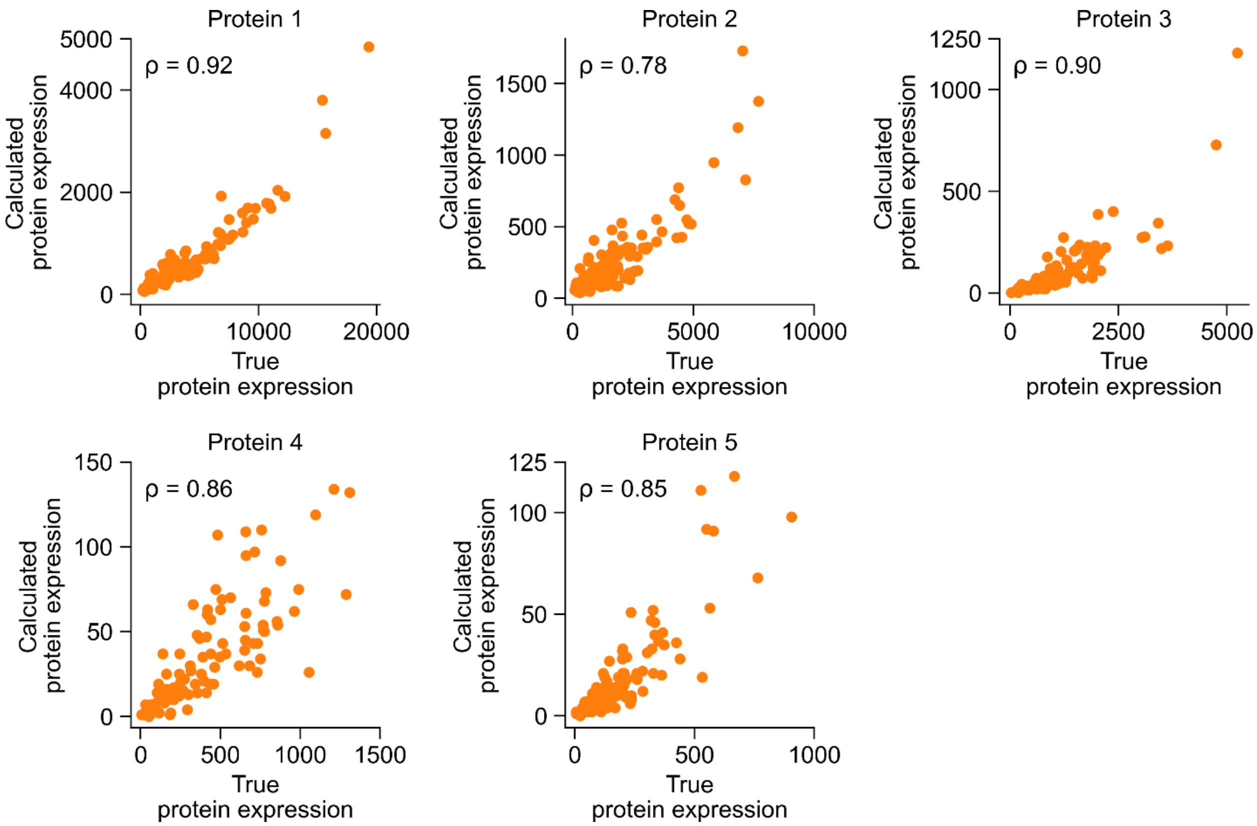


**Figure S4. Comparison between true and calculated protein expression.** Scatter plots showing the correlation between true and calculated protein expression in simulated data. The protein expression is equal to the UMI count of each protein per single cell. Each panel also displays the corresponding Spearman's correlation coefficient, ρ.

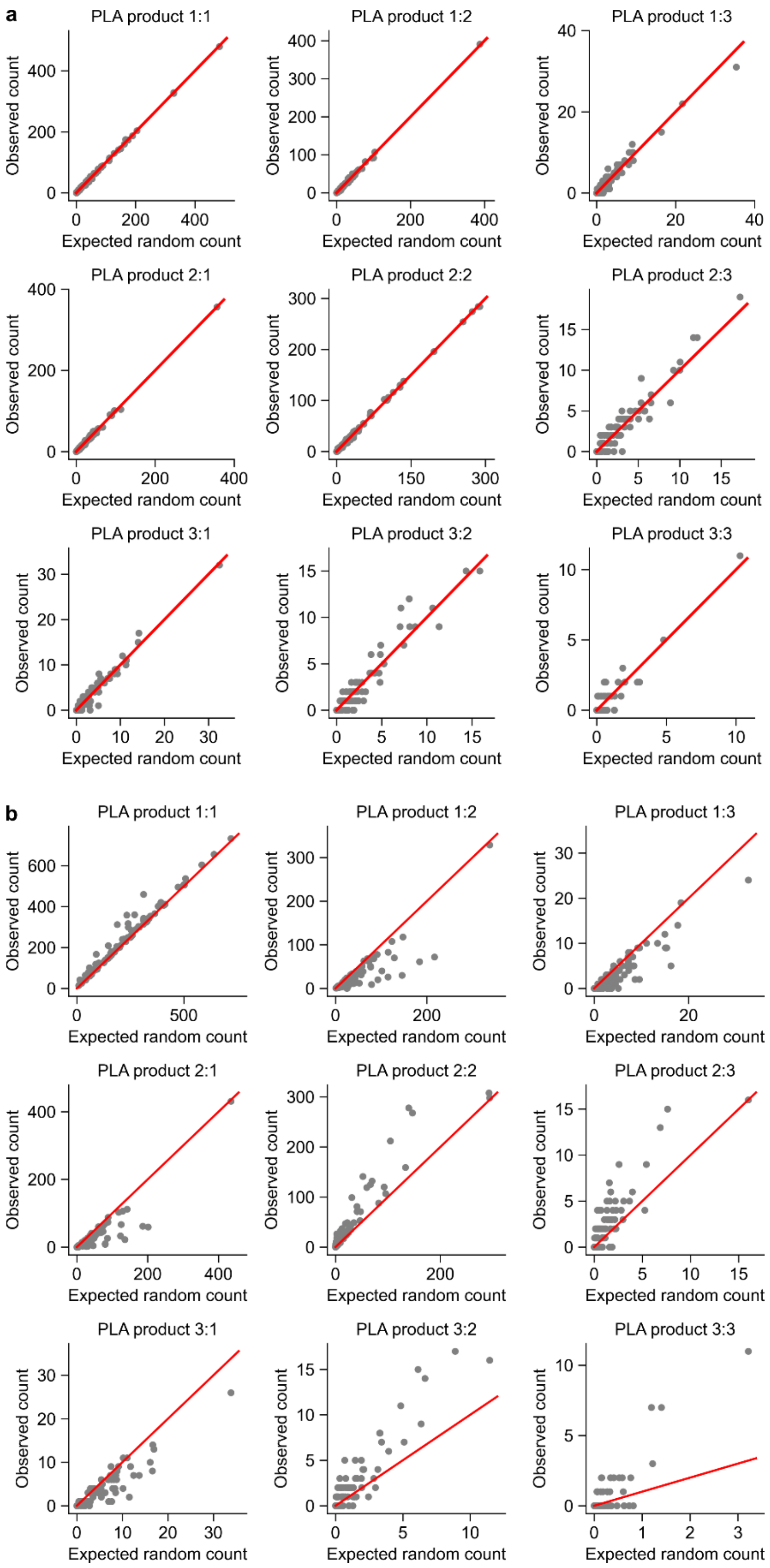


**Figure S5. Comparison between observed and expected PLA product count in simulated data**. (a, b) Scatter plots showing the observed and expected random count of each PLA product in the scenario when (a) no protein complex, and when (b) 1:1 is the only protein complex. The red lines indicate y = x.

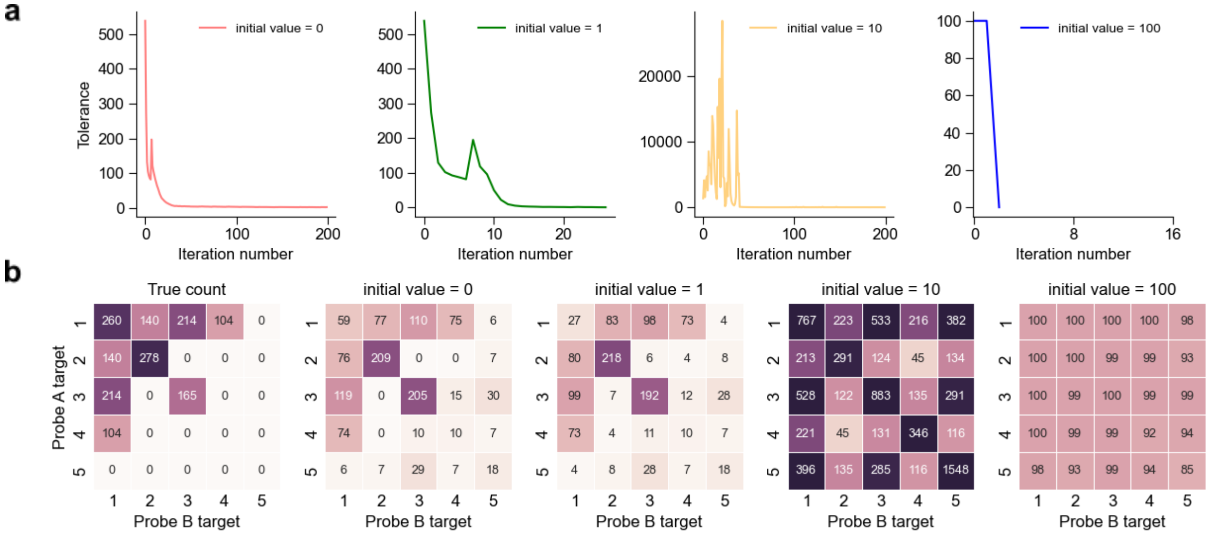


**Figure S6. The iterative method is sensitive to initial values.** (a) The change in output for each iteration (tolerance) changes depending on the initial values given to the algorithm. (b) The resulting protein complex estimates for each initialization, compared to the true complex values (left panel).

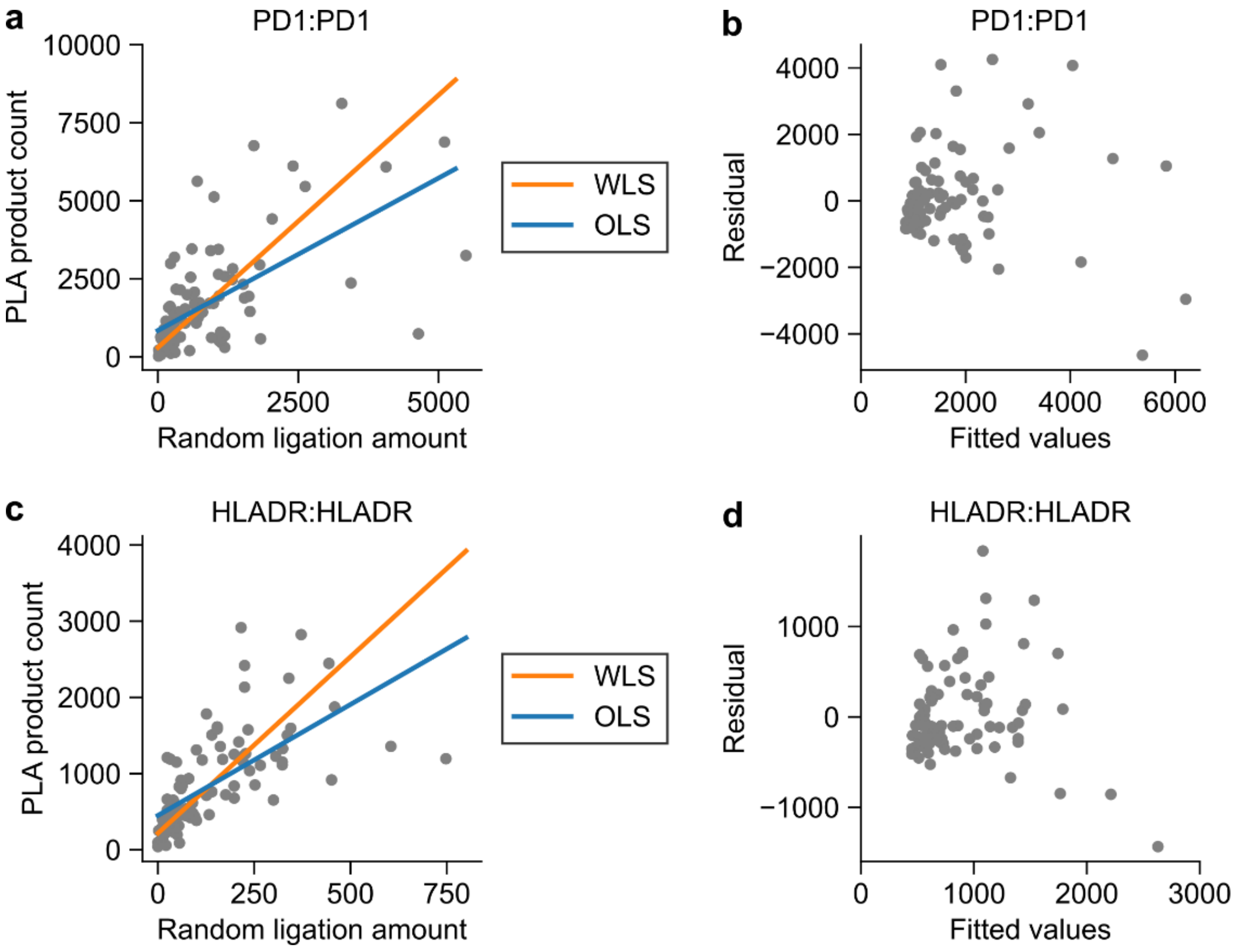


**Figure S7. Heteroscedasticity in PLA product count.** (a) Scatter plot showing the relationship between observed count of PLA product PD1:PD1 and the measured random ligation amount in Jurkat cells, and the corresponding weighted least squares (WLS) and ordinary least squares (OLS) regression lines. (b) Residual plot of ordinary least squares regression for PLA product PD1:PD1 in Jurkat cells. (c) Scatter plot showing the relationship between observed count of PLA product HLADR:HLADR and the measured random ligation amount in Raji cells, and the corresponding WLS and OLS regression lines. (d) Residual plot of ordinary least squares regression for PLA product HLADR:HLADR in Raji cells.

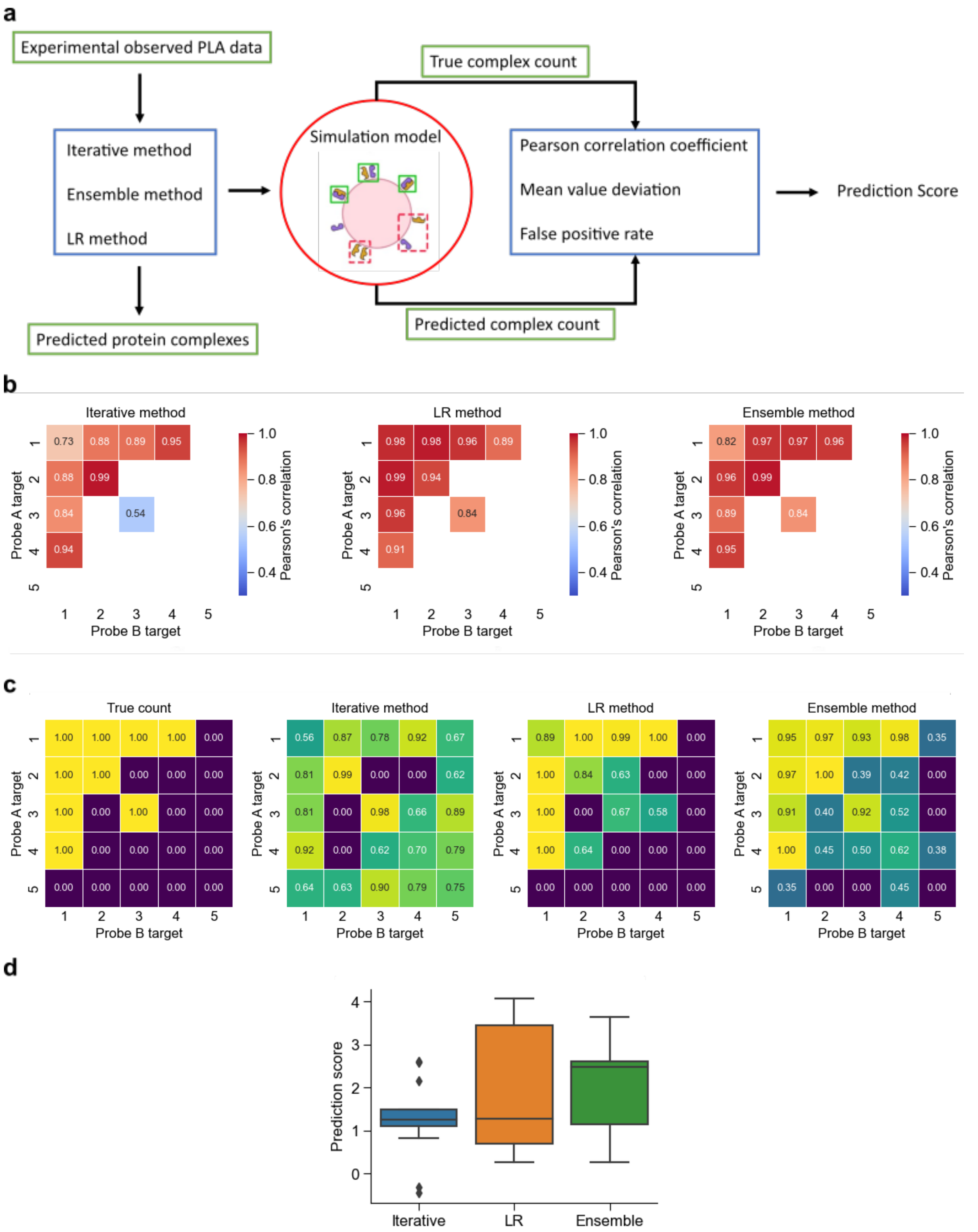


**Figure S8. Overview of the prediction score**. (a) Scheme of using simulation model to evaluate prediction algorithms with a quantitative scoring strategy. (b) The Pearson's correlation coefficient between true and predicted complex counts for each method. (c) The percent of cells called to have a protein complex for each method, along with the true counts. (d) The median prediction score of the ensemble method is higher than both the LR and iterative when comparing across all scenarios. Ensemble also displays a lower variance than LR.

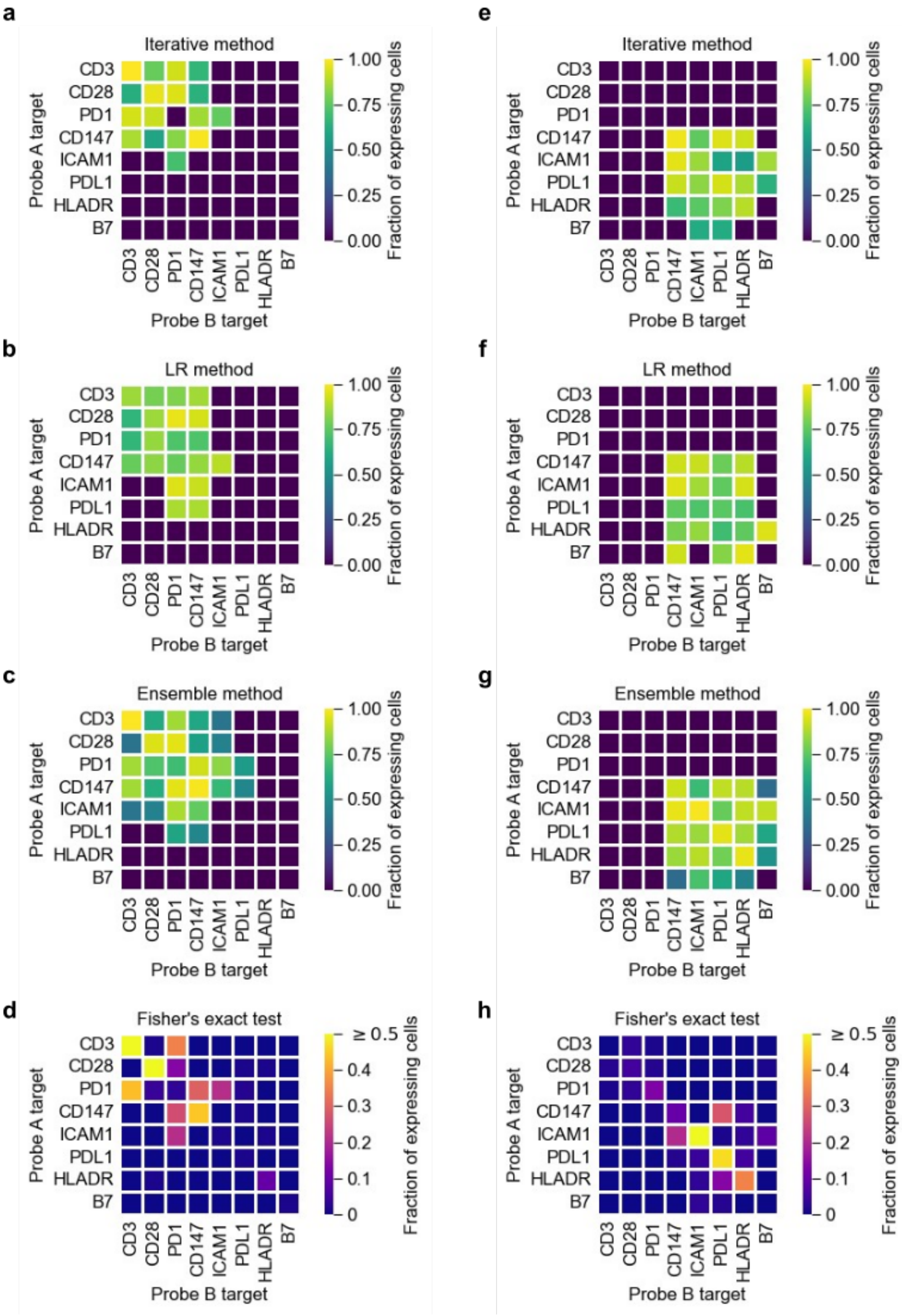
a
Iterative method
b
LR method
c
Ensemble method
d
Fisher's exact test
e
Iterative method
f
LR method
g
Ensemble method
h
Fisher's exact test
Probe A target
Probe B target
CD3
CD28
PD1
CD147
ICAM1
PDL1
HLADR
B7
Fraction of expressing cells
1.00
0.75
0.50
0.25
0.00
≥ 0.5
0.4
0.3
0.2
0.1
0

**Figure S9. Comparison between the iterative method, the LR method, Ensemble, and Fisher's exact test for protein complex detection in real data**. (a-d) Heatmaps showing the fraction of Jurkat cells that express a protein complex, as predicted by (a) the iterative method, (b) the LR method, (c) the Ensemble method, and (d) the Fisher's exact test. (e-h) Heatmaps showing the fraction of Raji cells that express a protein complex, as predicted by (e) the iterative method, (f) the LR method, (g) the Ensemble method, and (h) the Fisher's exact test.

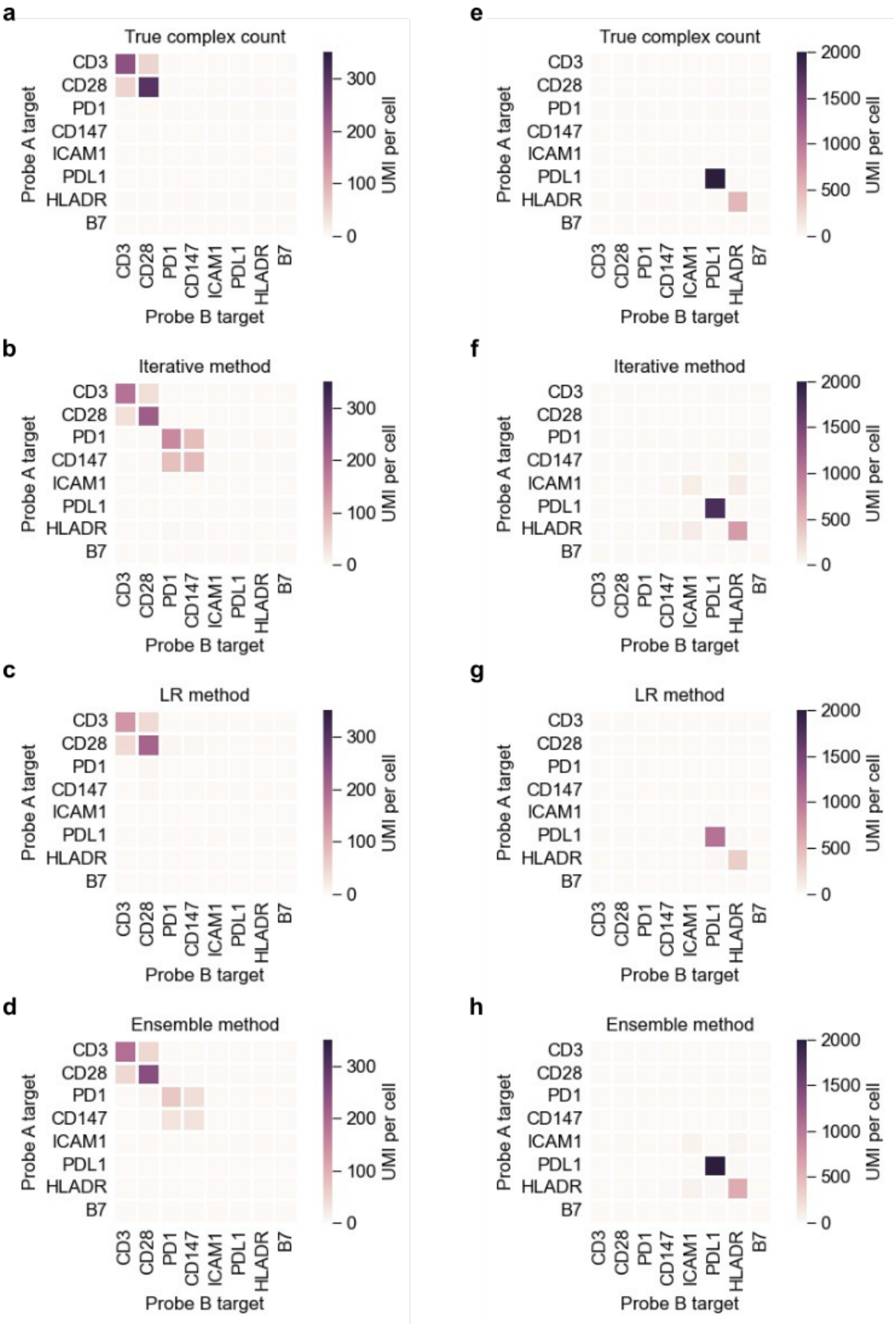
a
True complex count
e
True complex count
b
Iterative method
f
Iterative method
c
LR method
g
LR method
d
Ensemble method
h
Ensemble method
Probe A target
Probe B target
CD3
CD28
PD1
CD147
ICAM1
PDL1
HLADR
B7
UMI per cell
0
100
200
300
500
1000
1500
2000

**Figure S10. Simulation of T cells' and B cells' experimental data.** (a-d) Heatmaps of (a) true protein complex counts, and protein complex counts predicted by (b) iterative method, (c) LR method, and (d) Ensemble method in T cell simulation. (e-h) Heatmaps of (e) true protein complex counts, and protein complex counts predicted by (f) iterative method (g) LR method, and (h) the Ensemble method in B cell simulation. Here, the simulation parameters were chosen such that the total protein abundance was similar to experimental data. T cells were simulated to only express the protein complexes CD3:CD3, CD28:CD28, CD3:CD28 and CD28:CD3. B cells were simulated to only express the protein complexes PDL1:PDL1 and HLADR:HLADR.

### Supplementary tables

**Table S1.** List of simulation parameters.

| Parameter | Values | Note |
|---|---|---|
| R<br>$d_{ligation}$ | 5000<br>50 | Cell radius (1 unit = 1 nm)<br>Ligation distance |
| Figures S1, S3, S4<br>$c_{i,j}$<br>$A_i$<br>Bj | $\begin{bmatrix} 100 & 50 & 0 & 0 & 0 \\ 50 & 0 & 0 & 0 & 0 \\ 0 & 0 & 0 & 0 & 0 \\ 0 & 0 & 0 & 0 & 0 \\ 0 & 0 & 0 & 0 & 0 \end{bmatrix}$<br>$[2000 \quad 1000 \quad 500 \quad 200 \quad 100]$<br>$[2000 \quad 1000 \quad 500 \quad 200 \quad 100]$ | True protein complex count<br>Non-interacting probe A count<br>Non-interacting probe B count |
| Figure 3<br>$c_{i,j}$<br>$A_i$<br>Bj | $\begin{bmatrix} 850 & 750 & 0 \\ 750 & 1400 & 0 \\ 0 & 0 & 0 \end{bmatrix}$<br>$[20 \quad 15 \quad 2]$<br>$[20 \quad 15 \quad 2]$ | True protein complex count<br>Non-interacting probe A count<br>Non-interacting probe B count |
| Figure 4b<br>$c_{i,j}$<br>$A_i$<br>Bj | $\begin{bmatrix} 3000 & 1500 & 2000 & 1000 & 0 \\ 1500 & 3000 & 0 & 0 & 0 \\ 2000 & 0 & 2000 & 0 & 0 \\ 1000 & 0 & 0 & 0 & 0 \\ 0 & 0 & 0 & 0 & 0 \end{bmatrix}$<br>$[30 \quad 30 \quad 20 \quad 10 \quad 10]$<br>$[30 \quad 30 \quad 20 \quad 10 \quad 10]$ | True protein complex count<br>Non-interacting probe A count<br>Non-interacting probe B count |
| Figure 4c<br>$c_{i,j}$<br>$A_i$<br>Bj | $\begin{bmatrix} 300 & 150 & 200 & 100 & 0 \\ 150 & 300 & 0 & 0 & 0 \\ 200 & 0 & 200 & 0 & 0 \\ 100 & 0 & 0 & 0 & 0 \\ 0 & 0 & 0 & 0 & 0 \end{bmatrix}$<br>$[3000 \quad 1000 \quad 3000 \quad 1000 \quad 1000]$<br>$[3000 \quad 1000 \quad 3000 \quad 1000 \quad 1000]$ | True protein complex count<br>Non-interacting probe A count<br>Non-interacting probe B count |

| | | |
|---|---|---|
| Figure S2 | | Non-interacting probe A count |
| $c_{i,j}$ | $\begin{bmatrix} 0 & 0 & 0 \\ 0 & 0 & 0 \\ 0 & 0 & 0 \end{bmatrix}$ | Non-interacting probe B count |
| $A_i$ | $[2 \quad 20.1 \quad 0.6]$ | Non-interacting probe A count |
| Bj | $[5.7 \quad 41.6 \quad 0.1]$ | Non-interacting probe B count |
| Figure S5a | | |
| $c_{i,j}$ | $\begin{bmatrix} 0 & 0 & 0 \\ 0 & 0 & 0 \\ 0 & 0 & 0 \end{bmatrix}$ | True protein complex count |
| $A_i$ | $[1000 \quad 1000 \quad 100]$ | Non-interacting probe A count |
| Bj | $[1000 \quad 1000 \quad 100]$ | Non-interacting probe B count |
| Figure S5b | | |
| $c_{i,j}$ | $\begin{bmatrix} 200 & 0 & 0 \\ 0 & 0 & 0 \\ 0 & 0 & 0 \end{bmatrix}$ | True protein complex count |
| $A_i$ | $[1000 \quad 1000 \quad 100]$ | Non-interacting probe A count |
| Bj | $[1000 \quad 1000 \quad 100]$ | Non-interacting probe B count |
| Figure S9 | | |
| $c_{i,j}$ | $\begin{bmatrix} 240 & 50 & 0 & 0 & 0 & 0 & 0 & 0 \\ 50 & 300 & 0 & 0 & 0 & 0 & 0 & 0 \\ 0 & 0 & 0 & 0 & 0 & 0 & 0 & 0 \\ 0 & 0 & 0 & 0 & 0 & 0 & 0 & 0 \\ 0 & 0 & 0 & 0 & 0 & 0 & 0 & 0 \\ 0 & 0 & 0 & 0 & 0 & 0 & 0 & 0 \\ 0 & 0 & 0 & 0 & 0 & 0 & 0 & 0 \\ 0 & 0 & 0 & 0 & 0 & 0 & 0 & 0 \end{bmatrix}$ | True protein complex count |
| $A_i$ | $[50 \quad 330 \quad 3400 \quad 2400 \quad 16 \quad 3 \quad 100 \quad 1]$ | Non-interacting probe A count |
| Bj | $[50 \quad 330 \quad 3400 \quad 2400 \quad 16 \quad 3 \quad 100 \quad 1]$ | Non-interacting probe B count |
| Figure S10 | | |

| | | |
|---|---|---|
| $c_{i,j}$ | $\begin{bmatrix} 0 & 0 & 0 & 0 & 0 & 0 & 0 & 0 \\ 0 & 0 & 0 & 0 & 0 & 0 & 0 & 0 \\ 0 & 0 & 0 & 0 & 0 & 0 & 0 & 0 \\ 0 & 0 & 0 & 0 & 0 & 0 & 0 & 0 \\ 0 & 0 & 0 & 0 & 0 & 0 & 0 & 0 \\ 0 & 0 & 0 & 0 & 0 & 3000 & 0 & 0 \\ 0 & 0 & 0 & 0 & 0 & 0 & 500 & 0 \\ 0 & 0 & 0 & 0 & 0 & 0 & 0 & 0 \end{bmatrix}$ | True protein complex count |
| $A_i$ | $[5 \quad 60 \quad 30 \quad 1000 \quad 2500 \quad 3 \quad 5000 \quad 4]$ | Non-interacting probe A count |
| Bj | $[5 \quad 60 \quad 30 \quad 1000 \quad 2500 \quad 3 \quad 5000 \quad 4]$ | Non-interacting probe B count |

For simulation parameters of different biological scenario tests to generate prediction score table (Table S2), please refer to https://github.com/tay-lab/Prox-seq_computation.

**Table S2.** Prediction score under different simulation scenarios.

| Simulation contexts | | Iterative method | LR method | Ensemble method |
|---|---|---|---|---|
| Multiple protein complexes, 5 proteins | High signal, low noise | 2.15 | 0.82 | 2.54 |
| | High noise, low signal | 1.49 | 3.45 | 3.06 |
| Multiple protein complexes, 3 proteins | High signal, low noise | 1.28 | 4.07 | 3.57 |
| | High noise, low signal | 1.13 | 3.55 | 2.51 |
| | Similar signal & noise | 1.29 | 3.62 | 3.65 |
| Only homodimers, 3 proteins | High signal, low noise | 1.09 | 0.55 | 1.05 |
| | High noise, low signal | 1.24 | 1.18 | 1.14 |
| Only heterodimers, 3 proteins | High signal, low noise | 2.61 | 0.70 | 2.48 |
| | High noise, low signal | 2.59 | 2.54 | 2.60 |
| One over-abundant protein forming complexes, 3 proteins | High signal, low noise | 1.23 | 1.27 | 1.76 |
| | High noise, low signal | 0.83 | 2.08 | 2.08 |
| One over-abundant protein forming only | High signal, low noise | -0.32 | 0.27 | 0.27 |

| | | | | |
|---|---|---|---|---|
| homodimer, 3 proteins | High noise, low signal | -0.45 | 0.45 | 0.41 |
